# Plastic mulches reduce adult and larval populations of *Drosophila suzukii* in fall-bearing raspberry

**DOI:** 10.1101/2021.05.17.444501

**Authors:** Hanna McIntosh, Amaya Atucha, Philip A Townsend, W Beckett Hills, Christelle Guédot

## Abstract

The invasive spotted-wing drosophila, *Drosophila suzukii*, is a major pest of fruit crops world-wide. Management of *D. suzukii* relies heavily on chemical control in both organic and conventional systems, and there is a need to develop more sustainable management practices. We evaluated the efficacy of three colors of plastic mulches at reducing populations of *D. suzukii* in fall-bearing raspberry and assessed the mulches’ impacts on canopy microclimate factors relevant to *D. suzukii*. Black, white, and metallic plastic mulches reduced adult *D. suzukii* populations by 42-51% and larval populations by 52-72% compared to the grower standard. The mulches did not change canopy temperature or relative humidity, but metallic mulches increased canopy light intensity compared to the black mulch. Radiance in the visible spectrum (401-680 nm) was higher for the white and metallic mulch plots, but the black mulch plots did not differ from the control. In the UV spectrum (380-400 nm), all three plastic mulches had higher radiance than the control plots. Future studies will determine whether changes in radiance are associated with the observed reduction in *D. suzukii* populations. Plastic mulches are a promising cultural practice for managing *D. suzukii* since they can reduce adult and larval populations and could be incorporated into an integrated pest management program in both organic and conventional systems.

## Introduction

Production of small fruit crops is threatened by damage from spotted-wing drosophila, *Drosophila suzukii* (Diptera: Drosophilidae), an invasive pest of soft-skinned fruit crops. First detected in the continental US in 2008 (Hauser 2011), the fly has quickly spread from its native range in Eastern Asia into most major fruit-producing regions of the world (CABI 2016). Females lay their eggs in undamaged, ripening fruit using their serrated ovipositor (Kanzawa 1939; Walsh et al. 2011), and larvae feed inside the fruit, resulting in unmarketable fruit. Susceptible crops include raspberry, blackberry, blueberry, strawberry, sweet and tart cherry, and some cultivars of wine grapes (Lee et al. 2011; Bellamy et al. 2013; Ioriatti et al. 2015; Pelton et al. 2017; Kamiyama and Guedot 2019). For growers producing for fresh market or pick-your-own operations, *D. suzukii* damage substantially reduces the yield of marketable fruit, making susceptible crops difficult to grow sustainably (Farnsworth et al. 2017; DiGiacomo et al. 2019). For growers producing for fruit processing, *D. suzukii* infestation can lead to complete loss because of zero-tolerance policies for insect infestation set by processors (Bruck et al. 2011).

Currently, *D. suzukii* is managed primarily using chemical control in both organic and conventional systems (Haye et al. 2016). Recommendations for management include applying broad-spectrum insecticides every 4-7 days from detection of *D. suzukii* until harvest (Van Timmeren and Isaacs 2013). In California, the cost of chemical controls for *D. suzukii* was estimated at $1,161 per hectare for conventional and $2,933 for organic growers (Farnsworth et al. 2017). Few effective insecticides are approved for organic production (Sial et al. 2019), giving organic growers limited options for chemical control. Recently, *D. suzukii* has been reported to show reduced sensitivity to some active ingredients, including spinosad, the main insecticide used to control *D. suzukii* in organic systems (Van Timmeren et al. 2018; Gress and Zalom 2019). Other concerns about frequent insecticide use include negative health impacts on farm workers from pesticide exposure (McCauley et al. 2006; Flocks 2012; Schwartz et al. 2015), declining populations of beneficial insects (Roubos et al. 2014), and potential secondary pest outbreaks (Sarkar et al. 2020). Cultural practices can help manage *D. suzukii*, including cultivar selection, pruning or trellising, exclusion netting, harvesting fruit promptly, field sanitation, proper disposal of infested fruit, and post-harvest cold storage (Leach et al. 2016, 2018; Hooper and Grieshop 2020a, b; Schöneberg et al. 2020, 2021). However, these methods are labor intensive and often must be used in tandem with chemical control (Leach et al. 2016, 2018).

Cultural practices that modify the crop canopy microclimate have the potential to reduce *D. suzukii* infestation due to the fly’s sensitivity to temperature, humidity, and light (Tochen et al. 2016; Guédot et al. 2018). Adult summer morph *D. suzukii* thrive in warm but not hot temperatures, with the highest rate of population increase between 20-28 °C (Hamby et al. 2016) and an upper threshold for development at 30.9 °C (Ryan et al. 2016). Low populations of *D. suzukii* are observed in California in the summer, suggesting development or activity are slowed by hotter temperatures (Wang et al. 2016). *Drosophila suzukii* also thrives in very humid environments, with the longest survival and most eggs laid at 94% relative humidity in the lab (Tochen et al. 2016). In the field, females laid more eggs in the inner canopy of blackberry and blueberry, likely due to the darker, cooler, more humid environment (Diepenbrock and Burrack 2017; Evans et al. 2017). Thus, increasing the temperature and reducing humidity in the crop canopy could deter *D. suzukii* from laying eggs or disrupt immature development inside fruit.

Plastic mulches have been used in fruit and vegetable agroecosystems since the 1950s due to their ability to modify the microclimate, improving weed control, inducing earlier ripening, improving fruit quality, and increasing yield (Tarara 2000; Lamont 2005; Kasirajan and Ngouajio 2012). White and metallic plastic mulches reflect solar radiation and can increase canopy light intensity (Decoteau et al. 1989; Jakopic et al. 2007; Andreotti et al. 2010; Nottingham and Kuhar 2016; Smrke et al. 2019; Nottingham and Beers 2020) and canopy temperatures (Gordon et al. 2008; Andreotti et al. 2010) in fruit and vegetable crops. Despite the plastic mulches’ ability to modify the canopy microclimate, their efficacy as a cultural control for insect pests has been inconsistent by crop and insect species. For example, studies have reported that the use of black and white plastic mulches deterred aphids in watermelon and yellow squash (Farias-Larios and Orozco-Santos 1997; Greer and Dole 2003; Ban et al. 2009), but attracted aphids, whiteflies, and thrips in tomatoes (Greer and Dole 2003). Metallic plastic mulches reduced insect pest populations the most consistently, reducing Asian citrus psyllid (Croxton and Stansly 2014), tarnished plant bug (Rhainds et al. 2001), and pear psylla (Nottingham and Beers 2020). The use of plastic mulches could be an alternative strategy for managing *D. suzukii*, and no studies have yet evaluated their effectiveness. The only study evaluating the use of mulches to control *D. suzukii* tested a black fabric weedmat, which is commonly used for weed control. The results of this multi-state study were inconclusive, with only two locations reporting lower fruit infestation in plots with the black weedmat (Rendon et al. 2019).

The goal of this study was to evaluate the impact of black, white, and metallic plastic mulches on *D. suzukii* populations in fall-bearing raspberry. We assessed the impact of plastic mulches on adult and larval populations of *D. suzukii* and evaluated the ability of plastic mulches to modify canopy temperature, relative humidity, and light conditions. We expected that the white and metallic mulches would make the canopy microclimate less favorable to *D. suzukii*, thereby decreasing adult and larval populations. We did not expect the black mulch to impact the canopy microclimate and expected a smaller reduction in *D. suzukii* populations in plots with the black mulch.

## Materials and Methods

### Plot Establishment and Experimental Design

This study was conducted on a small commercial fruit and vegetable farm in Iowa County, WI, USA (42.99733N, -89.95843W) in 2019 and 2020. The raspberry plants were established in 2012 in Plano silt loam in rows 30 m long and 0.5 m wide with 3.05 m between rows for a total area of 0.08 hectares. Alley ways were planted with orchard grass and straw was applied in the rows during winter before canes emerged. The study was established in two rows each of fall-bearing cultivars ‘Caroline’ and ‘Polana’. Each plot was irrigated with 1.3 cm drip tape with emitters every 5.1 cm, which was placed down the middle of each row. Plots were irrigated identically when needed. Weeds were removed by hand from the gap between mulches and in the control plots as necessary. No insecticides were applied in 2019 or 2020.

The plastic mulch treatments included black biodegradable (Organix AG Film in 0.9 mil, Organix Solutions, Bloomington, MN), white-on-black biodegradable (Organix AG Film in 0.9 mil), and metallic-on-white polyethylene (SHINE N’RIPE in 1.25 mil, Imaflex, Montreal, Quebec), and a grower standard control, where grass filled in the space between the raspberry canes and the alleyway. The treatments were set up in a randomized complete block design with all four treatments in each row, totaling 16 treatment plots.

### Adult and Larval Populations

The raspberry field was monitored weekly for the presence of *D. suzukii* using three Scentry SWD traps (Scentry Biologicals, Billings, MT) with a drowning solution of 100 mL apple cider vinegar and one drop of unscented dish soap (Seventh Generation, Burlington, VT). Once *D. suzukii* was detected, the adult population was passively monitored in the experimental plots using one 15.25 cm^2^ clear sticky card (Alpha Scents, West Linn, OR) placed in the fruiting zone in the center of each plot. The sticky cards were adjusted vertically as the plants grew to account for changes in canopy height. The sticky cards were replaced every 7 d, and the number of *D. suzukii* caught on each card was recorded.

To assess larval infestation of fruit, 36 ripe fruits (∼100 g) were randomly collected from each plot every two weeks from August 19 to October 8, 2019. The salt flotation method (Dreves et al. 2014) was used to determine the number of larvae in half of the fruit sample. The other half was placed in plastic cups and flies were reared to the adult stage to determine the proportion of *D. suzukii* to other Drosopholids. Rearing cups were kept at ambient lab conditions for 3 weeks, and then all flies were identified. In 2020, sampling methods were modified slightly to allow for weekly sampling. Each week from August 25 to September 29, 2020, samples of 23 fruits were randomly collected from each plot and 18 fruits were used in the salt flotation tests and 5 fruits were placed in rearing cups. All flies that emerged from rearing cups in both years were identified as *D. suzukii*, so no adjustments were made to the data.

### Canopy Microclimate

Canopy temperature, relative humidity, and light intensity were monitored continuously from July 9 to October 18, 2019 and July 3 to October 6, 2020. HOBO data loggers were hung in the fruiting zone, and height was adjusted vertically as described for the sticky cards. Temperature and relative humidity were recorded every minute using HOBO U23 Pro v2 Data Loggers (OnSet, Bourne, MA) attached underneath a 25.4 cm diameter white plastic radiation shield. Light intensity was recorded every minute using HOBO Pendant MX Temperature/Light data loggers measuring in lux, a measure of the intensity of light between 400-700 nm, as perceived by the human eye.

We used a SVC HR 1024i spectrometer (Spectra Vista Corporation, Poughkeepsie, NY) to characterize radiance in the canopy. The 1024i measures radiance between 350-2500 nm across 1024 spectral channels (bands) with a resolution of 3.5 nm in the UV-visible, which is interpolated to 1 nm for analysis. Canopy light conditions were measured using a 25° field of view fiber optic cable with a pistol-grip for accurate targeting. Data in the treatment plots was collected in a single row of cultivar ‘Caroline’. On September 2-5, 2020, readings were taken in all four treatments on each day between 8 AM and 11 AM, when flies are active (Jaffe and Guédot 2019). The measurements were taken in a different sequence of treatments each day so that each treatment was measured first, second, third, and fourth to account for variation in light conditions by starting time. Radiance was measured at nadir (pointing straight downward) at 40 cm and 80 cm above the ground on both sides of the row at five evenly spaced horizontal positions along the plot. Three readings, each with a scan time of 2 seconds, were taken at every position for a total of 60 readings per treatment on each of the four measurement dates. Skies were clear on three of the four days and had consistent cloud cover on the other day. Outlier observations were identified and manually removed from analysis, but data from the cloudy day was not removed in order to characterize the radiance above the mulches in different sky conditions experienced by *D. suzukii*.

### Data Analysis

#### Adult and Larval Populations

Adult and larval populations were compared among treatments using a two-step approach analyzing zero and non-zero data separately to meet model assumptions (Cragg 1971). Logistic regressions were used to analyze the presence of adults or larvae with a binary indicator of *D. suzukii* presence / absence as the response variable. Linear mixed-effects models with log-transformed response variables were used to compare differences in the number of *D. suzukii* when present. For all models, the fixed effects were mulch treatment, row, and year, and the random effect was week crossed with year. In the model for the number of larvae in fruit samples, the female population trapped on sticky cards in the week before fruit sampling was included as an additional fixed effect. Tukey post-hoc tests were conducted following significance of fixed effects. To determine the overall change in adult and larval populations over the two years, percent change from controls was calculated. The two-year infestation rate was calculated for each mulch treatment [(total number of adults or larvae)/total number of days or fruits sampled], divided by the two-year infestation rate for the controls, and multiplied by 100. This number was subtracted from 100 for the percent reduction compared to the control.

#### Canopy Microclimate

Data for 2019 and 2020 were analyzed separately due to missing data from July to August 2019 caused by a data logger malfunction. Differences in daily mean, maximum, and minimum temperature and daily mean and minimum relative humidity were analyzed using generalized least squares regression with a moving average of order 3 (Box and Jenkins 1970; Koreisha and Pukkila 1990). The fixed effects were mulch treatment and row. Mean RH was a fixed effect for the temperature models and mean temperature was a fixed effect for the relative humidity models. Since most maximum relative humidity values were equal to 100%, the data was analyzed using a two-step approach. Logistic regressions were used to analyze occurrence of relative humidity readings equal to 100% with a binary indicator of 100% RH / <100% RH as the response variable. Ordinary least squares linear models were used to compare differences in maximum RH values <100%. The fixed effects were mulch treatment, row, and date.

Differences in the number of minutes per day above *D. suzukii’*s thermal developmental threshold of 30.9 °C (Ryan et al. 2016) and below 70% RH (as in Schöneberg et al. 2020) were analyzed separately using negative binomial regressions. For both models, the fixed effects were mulch treatment, row, and date.

Differences in mean and maximum daily light intensity were analyzed using ordinary least squares linear regression with log-transformed response variables. The fixed effects were mulch treatment, row, year, and a log-transformed one-day lag to account for autocorrelation of the data (Box and Jenkins 1970). Data from one experimental plot was removed from the 2020 data because logger placement was inconsistent with other plots due to small plant size.

The spectrometer data were calibrated to radiance using manufacturer-provided calibration coefficients and were further processed to correct for overlap regions between detectors using custom Python scripts (http://github.com/EnSpec/SpecDAL). Color vision is likely highly conserved in *Drosophila* species (Kelber and Henze 2013; Little et al. 2019), which are most sensitive to ultraviolet (UV), blue, and green light with less sensitivity of wavelengths above 600nm (Hardie 1979; Yamaguchi et al. 2010; Kelber and Henze 2013; Schnaitmann et al. 2013). However, Fountain et al. (2020) showed that D. suzukii is attracted to orange and red light (617 and 660 nm), so may be more sensitive to wavelengths above 600 nm than *D. melanogaster*. Our analyses of radiance therefore used wavelengths that plausibly may be visible to *D. suzukii*: 338-680 nm.

To identify whether the mulch treatments affected radiance at wavelengths visible to *D. suzukii*, we used an ordinary least squares model to compare treatment differences in the area below the radiance curve using the trapezoid method. The data was divided into UV (338-400 nm) or visible (401-680 nm) spectra using a binary indicator “spectrum” to identify treatment differences at each spectrum. The response variable was the log-transformed area below the curve of radiance. Fixed effects were treatment * spectrum interaction, position in the plot, and date. Tukey post-hoc tests were run to determine the differences among treatments for UV and visible spectra.

## Results

### Adult and Larval Populations

Fly populations differed for both years of the study, with higher populations observed in 2019 than 2020 (df: 1, χ2 = 11.36, P < 0.001). The mulch treatments reduced the presence of female flies (df = 3, χ2 = 10.27, P = 0.016), with females present less often on sticky cards in the black mulch plots than in the control plots. The mulch treatments also reduced the number of female flies (df = 3, χ2 = 19.82, p < 0.001; Fig. 2a), with lower numbers of females trapped in all mulch treatments compared to the controls. The mulch treatments reduced the presence of male flies (df = 3, χ2 = 20.64, P < 0.001), with males present less often on sticky cards in the black and white mulch plots compared to the control plots. There was no difference in the number of male flies trapped in any of the treatments (df = 3, χ2 = 6.59, p = 0.09; Fig. 2b). Over the duration of the study, the total number of flies trapped was reduced by 51% in the black and metallic mulch treatments and by 42% in the white mulch treatment compared to the control.

**Fig. 1.**
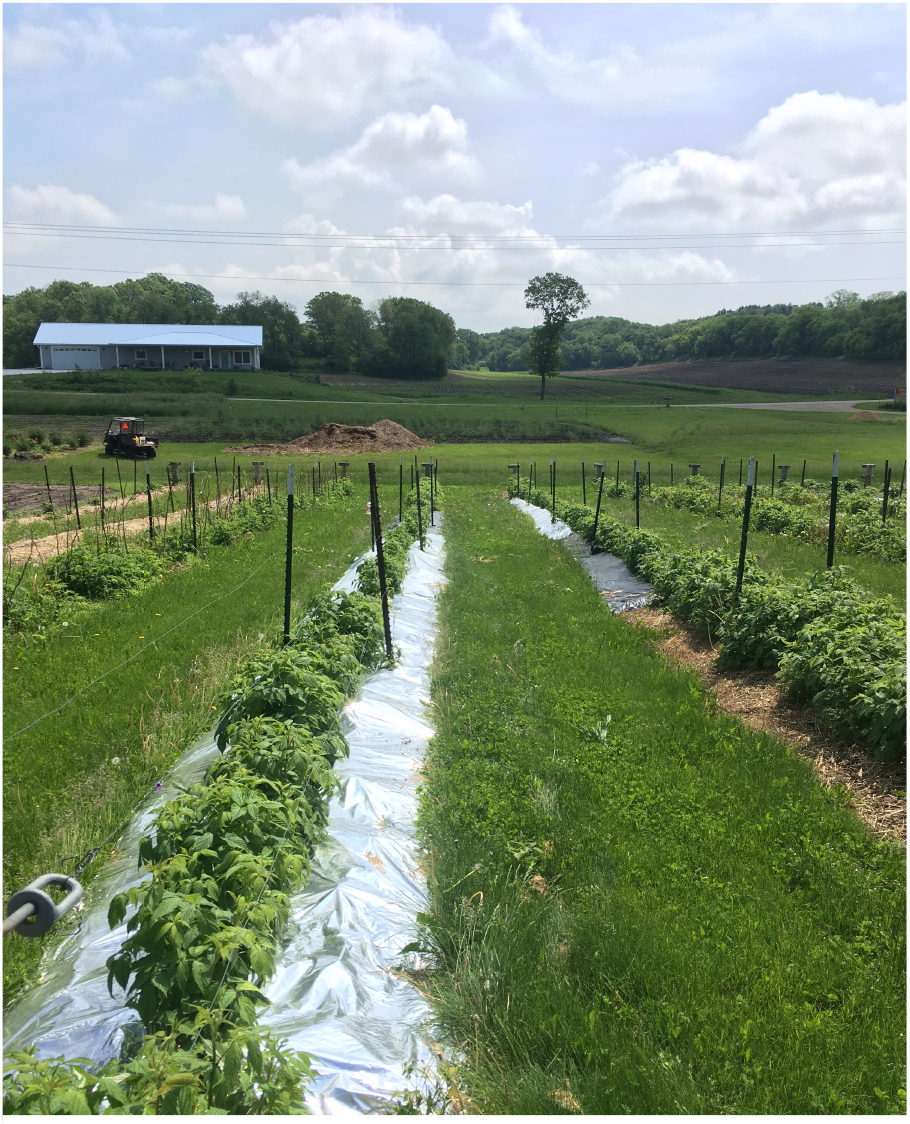
Plastic mulch strips laid along both sides of raspberry rows at grower-collaborator’s farm in Wisconsin Mulches were applied by hand in late April when raspberry canes were just emerging from below the straw mulch. Two mulch strips 7.6 m long and 70 cm wide were laid on each side of the row with a ∼10 cm gap down the center of the row for canes to grow (Fig. 1). All edges were secured with 15.25 cm biodegradable stakes (Eco Turf Midwest, Bensenville, IL) spaced ∼ 30 cm apart. Since the plastic mulches restricted the area where raspberry canes could emerge, canes in the control plots that grew outside the 10 cm center strip were pruned both years in June.

**Fig. 2.**
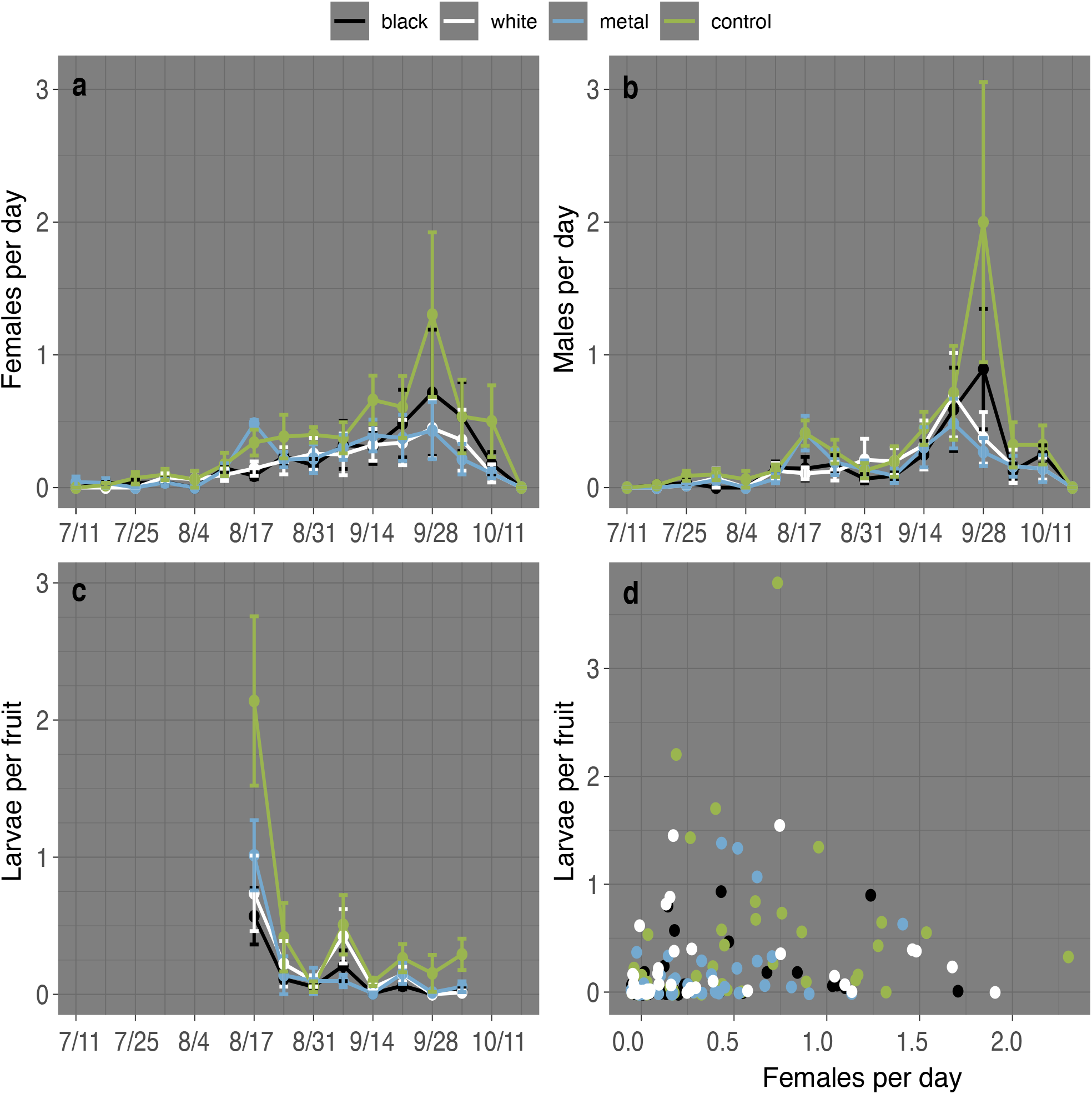
Effect of plastic mulches on adult and larval D. suzukii populations assessed in 2019 and 2020. Mean (± SE) number of (a) female and (b) male D. suzukii captured per day on clear sticky cards placed in the fruiting zone and (c) mean (± SE) number of larvae per fruit assessed using the salt float method. (d) Relationship between larval infestation of fruit and female population in the previous week

Larvae were present in fruit samples more often in 2019 than 2020 (df = 1, χ2 = 6.18, P = 0.013), but the number of larvae in fruit was not different between years (df = 1, χ2 = 2.09, P = 0.15). The mulch treatments reduced the presence of larvae in fruit samples (df = 3, χ2 = 8.02, P = 0.046), with larvae present marginally less in the black mulch plots compared to the control plots (p = 0.053). The mulch treatments also reduced the number of larvae in fruit samples (df = 3, χ2 = 24.42, p < 0.001; Fig. 2c), with fewer larvae in fruit from all the mulch treatment plots compared to the control plots. Over two years, the total number of larvae in sampled fruit was reduced 72% by the black mulch, 61% by the metallic mulch, and 52% by the white mulch. The female fly population in the week prior to fruit sampling was not a significant predictor of presence (df = 3, χ2 = 0.01, P = 0.92) or number (df = 3, χ2 = 0.54, P = 0.46; Fig. 2d) of larvae in fruit.

### Canopy Microclimate

There were no differences among treatments for daily mean, maximum, or minimum temperature or relative humidity in the raspberry canopy in 2019 or 2020 (Table S1, Table S2). There were also no differences among treatments in the number of minutes per day above the 30.9 °C developmental threshold or above 70% RH (Table S3). Year was a significant factor in both the mean and maximum light intensity models, with higher light intensity in 2019 than 2020. Mean light intensity was not impacted by the mulch treatments (df = 3, F = 0.31, P = 0.82; Fig. 3a,b), but maximum light intensity was higher in the canopy in the metallic mulch plots compared to the black mulch plots (df = 3, F = 3.10, P = 0.026; Fig. 3 c,d).

**Fig. 3.**
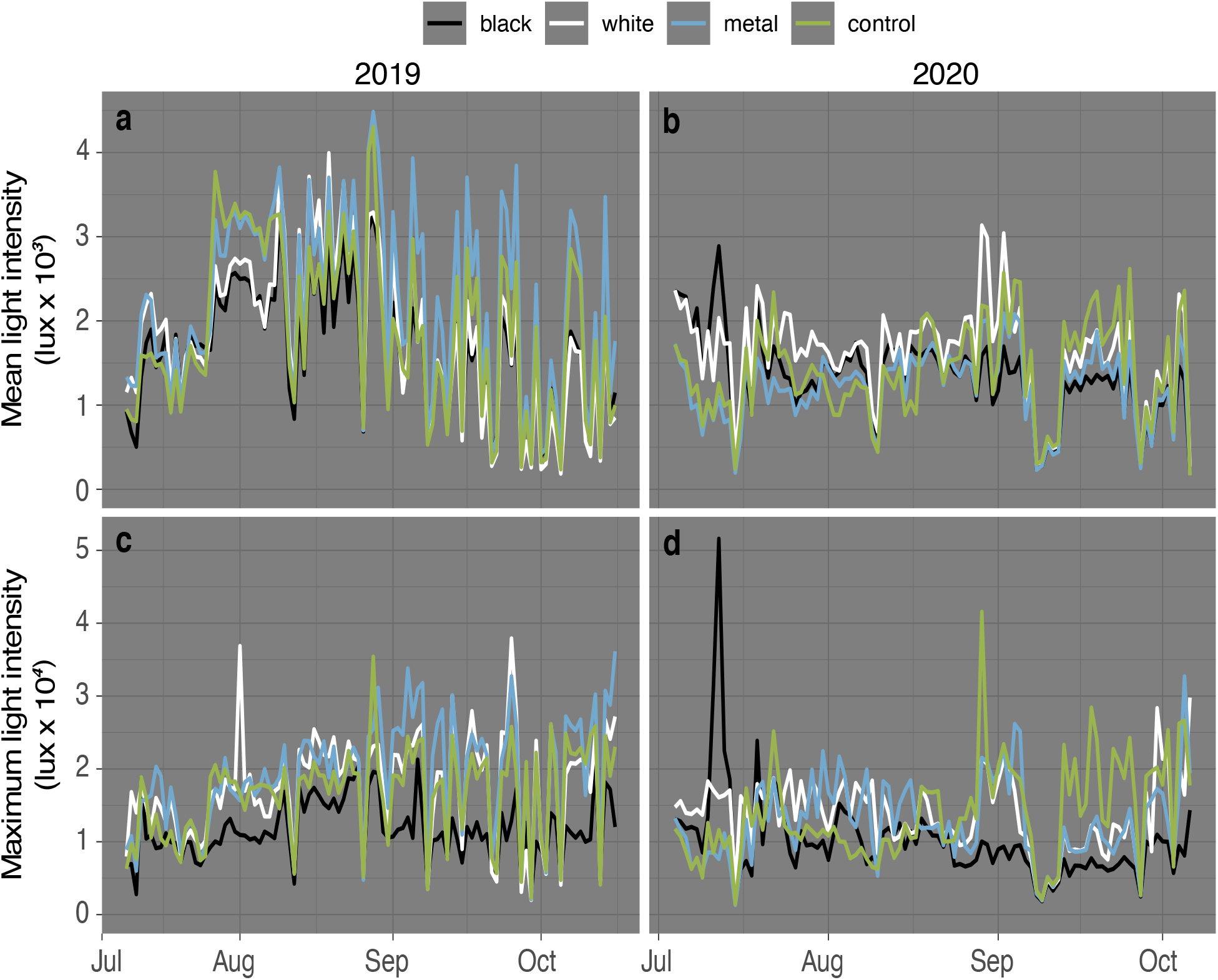
Effect of plastic mulches on (a,b) daily mean and (c,d) maximum light intensity in the canopy of raspberry plants in the 2019 and 2020 growing seasons. Data was recorded using HOBO sensors placed in the fruiting zone. Error bars were omitted for clarity

The mulch treatment * spectrum interaction had a significant effect on radiance (df = 3, F = 7.95, P < 0.001; Fig. 4). In the UV spectrum (338-400 nm), the control plots had the lowest radiance, black and white mulches were equivalently intermediate, and the metallic mulch had the highest radiance. In the visible spectra (401-680nm), the control plots and black mulch were equivalent, the white mulch had substantially higher radiance, and the metallic mulch had the highest radiance.

**Fig. 4.**
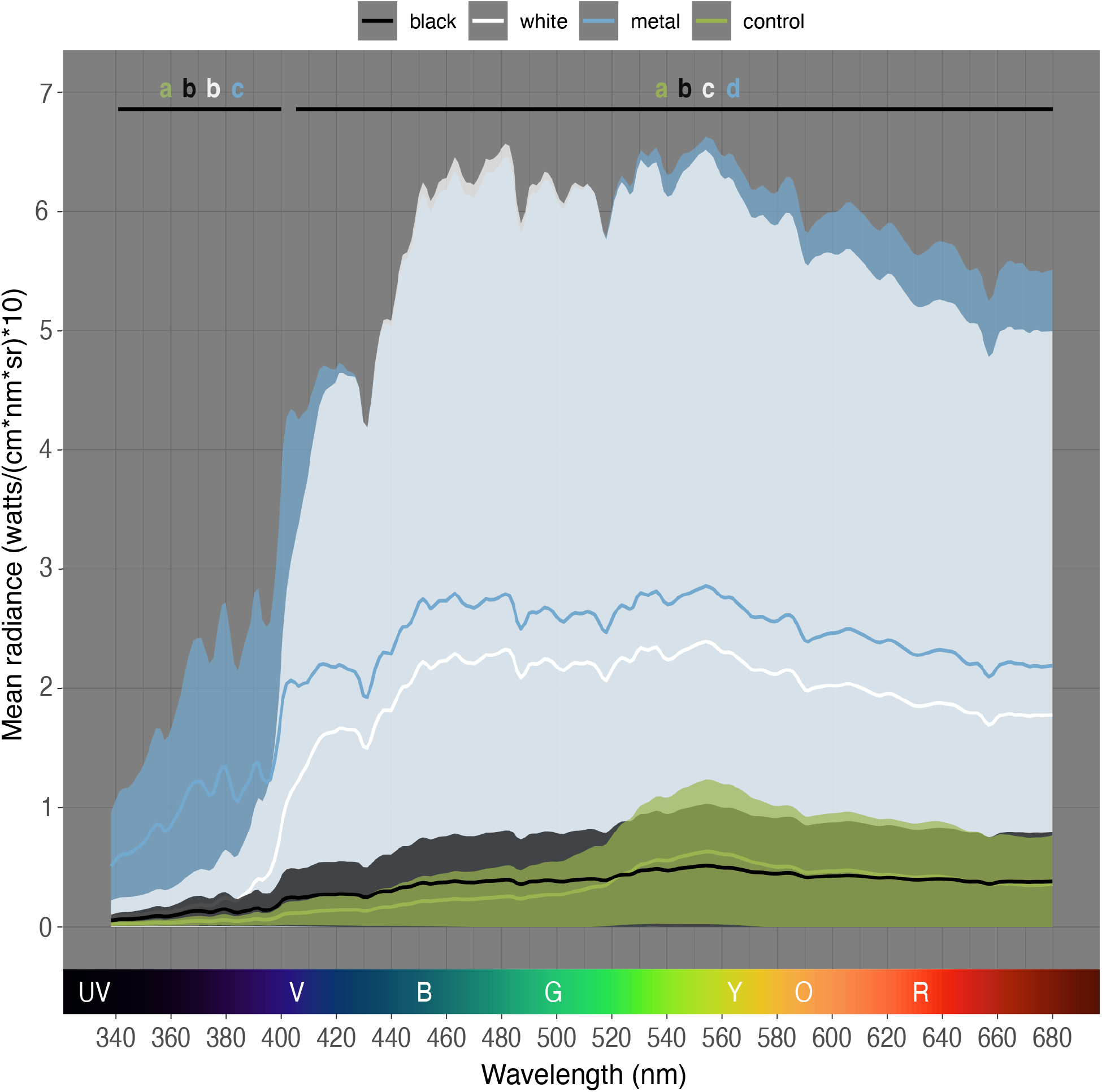
Mean (± SD) radiance in the canopy of raspberry plots with plastic mulch treatments. Treatments with different letters are significantly different from each other based on Tukey post-hoc tests

## Discussion

Black, white, and metallic plastic mulches reduced the number of adult female *D. suzukii* in the canopy and decreased larval infestation of raspberry fruits compared to the control. There were no differences in canopy temperature and humidity among treatments, but the maximum light intensity was higher for the metallic mulch than the black mulch. Radiance in the UV spectrum was higher in the canopy above all three plastic mulches compared to the control, while radiance in the visible spectrum was higher above the white and metallic mulches compared to the black mulch and control.

Our results show that black, white, and metallic plastic mulches reduced adult *D. suzukii* populations in the raspberry canopy by 42-51%, with the lowest adult populations in the black and metallic mulch plots. Similarly, other studies found that plastic mulches reduced pear psylla populations (Nottingham and Beers 2020), Asian citrus psyllids (Croxton and Stansly 2014) and Mexican bean beetles (Nottingham and Kuhar 2016). While there is clear evidence that plastic mulches can reduce adult insect populations, most studies did not separate populations by sex. We found that plastic mulches only affected populations of female *D. suzukii*, which is consistent with a study testing metallic plastic mulches for controlling Asian citrus psyllid (Ali and Qureshi 2020). Results from these two studies warrant further investigation of the sex-dependent effects of plastic mulches and provide insights into potential mechanisms by which plastic mulches reduce insect pest populations.

Plastic mulches tested in our study reduced adult female fly populations and also reduced larval infestation of fruit by 52-72% (Fig. 2), which is comparable to results from commonly used cultural practices like frequent harvesting and field sanitation. Harvesting fruit every 1-2 days reduced larval infestation by around 38-100% compared to a 3 day harvesting schedule (Leach et al. 2018). Composting cull fruit with chicken manure, burying it at a depth of 24 cm, or solarizing infested berries in plastic bags killed 95%, 97%, and 99% of *D. suzukii* larvae, respectively (Leach et al. 2018; Hooper and Grieshop 2020a, b). Practices that exclude *D. suzukii* from crops were similarly effective. Covering high tunnels with exclusion netting or plastic reduced *D. suzukii* larval infestation by 74% and 98%, respectively (Leach et al. 2016; Rogers et al. 2016). In contrast, plastic mulches appear to be more effective at reducing *D. suzukii* larval infestation than other cultural practices that modify the crop microclimate. In a multi-state study, woven black weedmat had no overall effect on *D. suzukii* emergence from blueberries (Rendon et al. 2019). Using drip irrigation instead of overhead irrigation did not reduce larval infestation of fruit and the use of overhead irrigation did not deter adult flies (Rendon and Walton 2019). Pruning approximately 50% of the raspberry canopy reduced *D. suzukii* larval infestation by about 13% (Schöneberg et al. 2020). Understanding how plastic mulches impact the canopy microclimate is thus very important as it may reveal why they are more effective than other microclimate modifying cultural practices.

Temperature and humidity have been shown to impact *D. suzukii* populations (Hamby et al. 2016; Guédot et al. 2018) and other studies have reported that plastic mulches can alter canopy microclimate (Gordon et al. 2008; Andreotti et al. 2010; Strik et al. 2020), so we had hypothesized that increased canopy temperature and reduced humidity would be the mechanism by which plastic mulches reduced *D. suzukii* populations. However, no differences were detected in canopy temperature and relative humidity among treatments. Our results are consistent with studies in snap beans (Nottingham and Kuhar 2016) and pears (Nottingham and Beers 2020), which found that black, white, and metallic mulches did not impact canopy temperature or humidity. In our study, canopy conditions in all treatments were frequently in the optimal range for *D. suzukii* (Hamby et al. 2016; Tochen et al. 2016), with temperatures between 20-28 °C and RH above 94% occurring almost daily. Temperatures infrequently exceeded 30.9 °C, the upper developmental threshold for *D. suzukii* (Ryan et al. 2016), which is consistent with a previous study in Wisconsin raspberries (Guédot et al. 2018). It is possible that our sensors were located too far from the mulches to detect differences in canopy temperature and humidity, as they were placed in the top of the canopy in the fruiting zone. Taking measurements closer to the ground, where most *D. suzukii* are found in raspberry during the morning and mid-afternoon (Jaffe and Guédot 2019) may help detect differences among treatments.

Although canopy temperature and relative humidity did not differ among treatments, the metallic mulch increased maximum light intensity (400-700 nm) in the canopy compared to the control (Fig. 3C,D). Nottingham and Kuhar (2016) also measured higher reflected light intensity above white and metallic mulches in snap beans, attributing the reported reduction of Mexican bean beetles to the change in light conditions. However, in our study, light intensity was higher only in the metallic mulch treatment, while the other mulches did not differ from the control. Thus, light intensity alone does not explain the reduction in *D. suzukii* adult and larval populations observed in all plastic mulch treatments.

The radiance data gives a more in-depth view of light conditions in the canopy, particularly in the UV (338-400 nm) versus visible (401-680 nm) spectra. In the visible spectrum, radiance was higher for the white and metallic mulch plots, but the black mulch plots did not have higher radiance than the control plots (Fig. 4). Similarly, in pear orchards, PAR (400-700 nm) was also highest in metallic mulch plots, and PAR values were not different among black mulch and control plots (Nottingham and Beers 2020). While female *D. suzukii* are attracted to orange and red light (617-660 nm), possibly as a mechanism for detecting fruit (Fountain et al. 2020), attraction to these wavelengths alone does not provide a clear explanation for the reduction in *D. suzukii* observed in all of the plastic mulch treatments.

In the UV spectrum, all three plastic mulches had higher radiance than the control plots (Fig. 4), suggesting that the increase in reflection of UV light could be involved in deterring adult female *D. suzukii*. Similarly, black and metallic mulches had higher UV light intensity than control plots in pear orchards, possibly leading to the reported deterrence of pear psylla in mulched plots (Nottingham and Beers 2020). There are multiple mechanisms by which increased UV light could be deterring *D. suzukii*, since insects can have complex visual responses to this spectrum. Many insects use UV light for navigation, perceiving patches of bright UV light as a reference for open sky when flying through vegetation (Cronin and Bok 2016). Larval infestation was lower in strawberry tunnels where UV light was blocked by cladding, suggesting that the presence of UV light is important for *D. suzukii* (Fountain et al. 2020). In contrast, our data suggests that increased UV light was associated with deterrence of *D. suzukii* females in the raspberry canopy. We measured the highest radiance in the lower canopy (40 cm above the mulch), so it is possible that this bright source of UV light near the ground is attracting *D. suzukii* away from the fruit in the canopy. Alternatively, *Drosophila* can orient relative to the sky using light polarization, the predictable polarization of light produced by sunlight or reflective surfaces (Weir and Dickinson 2012; Cronin and Bok 2016). Light polarization is detected by *Drosophila* in the UV range (Weir and Dickinson 2012), so reflection of UV light by the plastic mulches could simulate a strong source of ‘sunlight’ coming from the ground, potentially disrupting orientation and navigation. Croxton & Stansly (2014) also hypothesized that disruption of polarization vision by metallic mulch was disorienting Asian citrus psyllids and reducing populations in orange trees. However, this mechanism is likely to be sex-independent, so it does not explain why *D. suzukii* males were not impacted by the plastic mulches.

A possible explanation of the sex-dependent deterrence of *D. suzukii* is that UV light could be deterring female flies from laying eggs in fruit. While female *D. suzukii* were attracted to UV light in the laboratory (Fountain et al. 2020), mating status may be an important factor in phototaxy in UV light. In a study with *D. melanogaster* in the lab, only virgin females were attracted to UV light, and mated females preferred to lay eggs in dark substrate (Zhu et al. 2014). While this has not been directly tested for *D. suzukii*, it is consistent with the fly’s preference to lay eggs in the dark inner canopy (Diepenbrock and Burrack 2017; Evans et al. 2017). If *D. suzukii* females also prefer to lay eggs away from UV light, higher radiance in the UV spectrum could explain why the plastic mulches only reduced populations of female flies in the canopy. Our results show that the plastic mulches alter radiance in the raspberry canopy compared to the control plots, and future studies should investigate whether the increase in reflection of UV light by the plastic mulches is the mechanism reducing *D. suzukii* adult female populations.

While our study showed plastic mulches reduced *D. suzukii* adult female and larval populations, and our data suggests that UV light might be deterring the female flies, we did not directly address whether the mulches are 1) reducing larval infestation due to reduced adult female populations, 2) altering oviposition behavior to reduce larval infestation, or 3) increasing mortality of immature flies inside the fruit. First, in this study, the female fly population in the week prior to fruit sampling was not a predictor of larval infestation of fruit, suggesting that reduced larval infestation may not be solely a product of reduced female populations. These results corroborate findings from another study using baited traps in cold climate wine grapes (Pelton et al. 2017). In California raspberries, adult trapping of *D. suzukii* and larval infestation were generally correlated, but with frequent instances of low trap captures with high larval counts, suggesting that adult populations are not always correlated with larval infestation (Hamby et al. 2014). Second, plastic mulches could alter female *D. suzukii* oviposition behavior by deterring oviposition via increased reflection of UV light (as discussed previously) or by increasing fruit temperature. Female *D. suzukii* prefer to oviposit in fruit between 20-25 °C, with decreasing oviposition in fruit above 28°C and no oviposition above 35°C (Zerulla et al. 2017). While canopy temperatures were not elevated by the plastic mulches, it remains unknown how fruit temperature was impacted. Third, this study did not measure mortality of immature *D. suzukii* inside fruit, and elevated fruit temperature could also increase mortality of eggs or larvae if fruit temperature exceeds the developmental threshold for *D. suzukii* (Ryan et al. 2016). Future studies should address these remaining gaps in knowledge.

This study demonstrates that plastic mulches are an effective cultural practice for managing *D. suzukii* in fall-bearing raspberry in the Upper Midwest, reducing adult and larval population by up to 51% and 72%, respectively. Applying plastic mulches is a preventative practice that could be used alongside other cultural controls, such as frequent harvesting and field sanitation, for a more robust integrated pest management program. The efficacy of plastic mulches seems to be influenced by agroecosystem, crop, and climate, so plastic mulches should be tested for *D. suzukii* management in other susceptible crops and climates to confirm efficacy in different regions. Overall, plastic mulches are a promising new tool that could be integrated in management programs for *D. suzukii* in both organic and conventional berry production systems.

## Declarations

### Authors’ contributions

HM, CG, and AA conceived and designed the study. Material preparation, data collection and analysis were performed by HM with advising from all other authors. The manuscript was written by HM, CG, and AA and all authors commented on previous versions of the manuscript. All authors read and approved the final manuscript.

## Acknowledgements

We acknowledge that this research was conducted on land forcibly taken from the Ho-Chunk people, and that the University of Wisconsin – Madison has built wealth in part from land seized from Indigenous people under the Morill Act. We recognize that agricultural and entomological research centers Western science and undervalues Indigenous knowledge, even though many sustainable and agroecological pest management tactics are Indigenous practices. We challenge ourselves and other researchers to continually evaluate how our research and outreach programs perpetuate colonialism and benefit from stolen land, and how we can decolonize our science by centering social justice, Indigenous knowledge and people, and land sovereignty.

We are grateful to our grower-collaborators Ed and Kathy Bures, and our grower advisory panel members Mike Koeppl, Andy Merry, Craig Carpenter, Mike Matushak, and Annie Deutsch. We thank Andi Nelson, Rodney Denu, Janet Hedcke, Matt Kamiyama, Matt Hetherington, Nolan Amon, and Bonnie Ohler for field assistance. We are grateful to Beth Workmaster, Erin Hokanson Wagner, and Ben Spaier for technical assistance. We thank Maria Kamenetsky for statistical consulting. Mulches for this project were generously donated by Organix Solutions and Imaflex.

