## Supplemental Tables for "Plastic mulches reduce adult and larval populations of *Drosophila suzukii* in fall-bearing raspberry"

|  |  |  | Mean temperature | | Minimum temperature | | Maximum temperature | |
| --- | --- | --- | --- | --- | --- | --- | --- | --- |
|  | Effect | df | LRT | p>χ2 | LRT | p>χ2 | LRT | p>χ2 |
| 2019 | Mulch | 3 | 0.009 | 1.00 | 0.00 | 1.00 | 0.23 | 0.97 |
|  | Row | 3 | 0.033 | 1.00 | 0.19 | 0.98 | 0.14 | 0.99 |
|  | RH | 1 | 2.61 | 0.11 | 317.43 | <0.001 | 261.76 | <0.001 |
| 2020 | Mulch | 3 | 0.040 | 1.00 | 0.11 | 0.99 | 0.78 | 0.85 |
|  | Row | 3 | 0.14 | 0.99 | 0.054 | 1.00 | 1.20 | 0.75 |
|  | RH | 1 | 26.49 | <0.001 | 56.81 | <0.001 | 167.37 | <0.001 |

**Table S1** Statistical parameters of generalized least squares models with a moving average of order 3 using mulch treatment, row, and RH as predictors of canopy temperature measured every minute using HOBO data loggers

**Table S2** Statistical parameters of generalized least squares models with a moving average of order 3 using mulch treatment, row, and temperature as predictors of mean and maximum canopy relative humidity measured every minute using HOBO data loggers. Maximum relative humidity was analyzed with mulch treatment, row, and date as predictors using logistic regressions with a binary indicator of 100% RH / <100% and ordinary least squares linear models for values <100%

|  | | | Mean RH | | | Minimum RH | | |  | | Maximum RH  100% vs <100% | | Maximum RH  <100% | | |
| --- | --- | --- | --- | --- | --- | --- | --- | --- | --- | --- | --- | --- | --- | --- | --- |
|  | Effect | df | LRT | p>χ2 | LRT | | p>χ2 | Effect | | df | χ2 | p>χ2 | | F | p>F |
| 2019 | Mulch | 3 | 0.38 | 0.94 | 0.64 | | 0.89 | Mulch | | 3 | 0.703 | 0.87 | | 0.44 | 0.72 |
|  | Row | 3 | 0.85 | 0.84 | 0.28 | | 0.96 | Row | | 3 | 1.60 | 0.66 | | 1.32 | 0.29 |
|  | Temp. | 1 | 30.99 | <0.001 | 30.47 | | <0.001 | Date | | 1 | 80.16 | <0.001 | | 9.05 | 0.006 |
| 2020 | Mulch | 3 | 0.40 | 0.94 | 0.70 | | 0.87 | Mulch | | 3 | 0.35 | 0.95 | | 0.45 | 0.72 |
|  | Row | 3 | 2.14 | 0.54 | 1.68 | | 0.64 | Row | | 3 | 1.47 | 0.69 | | 0.53 | 0.67 |
|  | Temp. | 1 | 88.65 | <0.001 | 4.96 | | 0.026 | Date | | 1 | 45.83 | <0.001 | | 11.06 | 0.002 |

**Table S3** Statistical parameters of negative binomial regressions using mulch treatment, row, and date as predictors of minutes per day above *D. suzukii’*s thermal developmental threshold of 30.9 ℃ and below 70% RH measured every minute using HOBO data loggers

|  |  |  | Minutes above 30.9 ℃ | | Minutes above 70% RH | |
| --- | --- | --- | --- | --- | --- | --- |
|  | Effect | df | F | p>F | χ2 | p>χ2 |
| 2019 | Mulch | 3 | 1.11 | 0.77 | 0.30 | 0.96 |
|  | Row | 3 | 0.31 | 0.96 | 1.79 | 0.62 |
|  | Date | 1 | 3.73 | 0.053 | 40.68 | <0.001 |
| 2020 | Mulch | 3 | 0.78 | 0.85 | 0.14 | 0.99 |
|  | Row | 3 | 0.28 | 0.96 | 2.92 | 0.40 |
|  | Date | 1 | 4.65 | 0.031 | 25.89 | <0.001 |
